# Nuclear and cellular, micro and nano calcification in Alzheimer’s disease patients and correlation to phosphorylated Tau

**DOI:** 10.1101/2020.06.15.148353

**Authors:** Elena Tsolaki, Lajos Csincsik, Jing Xue, Imre Lengyel, Sergio Bertazzo

## Abstract

Brain calcification (calcium phosphate mineral formation) has been reported in the past 100 years in the brains of Alzheimer’s disease (AD) patients. However, the association between calcification and AD, the triggers for calcification, and its role within the disease are not clear. On the other hand, hyperphosphorylated tau protein (pTau) tangles have been widely studied and recognized as an essential factor in developing AD. In this work, calcification in the brains of AD patients is characterized by advanced electron microscopy and fluorescence microscopy. Results are then compared to samples from cognitively healthy, age-matched donors, and the colocalization of calcification and pTau is investigated. Here, we show that AD patients’ brains present microcalcification associated with the neural cell nuclei and cell projections, and that these are strongly related to the presence of pTau. The link between microcalcification and pTau suggests a potential new mechanism of brain cell damage. Together with the formation of amyloid plaques and neurofibrillary tangles, microcalcification in neuronal cells adds to a better understanding of the pathology of AD. Finally, the presence of microcalcification in the neuronal cells of AD patients may assist in AD diagnosis, and may open new avenues for developing intervention strategies based on inhibition of calcification.

## 1. Introduction

Alzheimer’s disease (AD) is one of the most prevalent neurodegenerative diseases, affecting predominantly the elderly population[1]. It accounts for up to 80% of the cases of dementia and is responsible for a marked decline in cognitive abilities. One of the most well known hallmarks of the disease is the deposition of extracellular amyloid-β (Aβ)[2, 3] plaques and the intracellular neurofibrillary tangles (NFTs) formed from hyperphosphorylated tau proteins (pTau)[3, 4].

Tau protein is a microtubule protein primarily located in axons that, under normal conditions, interact with tubulin for microtubule stabilization[5]. This protein mainly functions through well-controlled phosphorylation events[6, 7], but under pathological conditions, it hyperphosphorylates[8] and form neurofibrillary tangles[9] which lead to the functional loss of neuronal cells. The development and the spread of tau pathology are recognized to be more relevant to brain regions associated with cognitive impairment[10].

AD has been previously associated with brain calcification (formation of calcium phosphate minerals)[11-18], generally in the form of intracranial vascular calcification in brain regions affected by the disease[19]. The origins, causes and role of this calcification are still not fully understood. Recently, a series of published works showed that an in-depth physicochemical characterisation of the calcification present in the most diverse diseases, such as aortic valve stenosis[20], atherosclerosis[20, 21], breast cancer[22] and macular degenerative diseases[23], may not only clarify the role of calcification, but also provide insight into the associated biological mechanisms of the disease itself. These methods, however, have never been applied to the characterization of the calcification present in AD patients. More importantly, there have been no studies to date, to the best of our knowledge, looking for possible correlations between the calcification physicochemical characteristics and the most relevant hallmarks of AD.

In this work, we analysed tissue sections from the basal ganglia, a region where brain calcification has been widely reported[11, 24-27], the temporal lobe, and the hippocampus, regions that have been strongly associated to AD[28-30]. Special attention was paid to the medial temporal lobe (middle, superior temporal and parahippocampal gyrus), a primary site of AD pathology[31-33]. Initially, the samples were characterized by Von Kossa staining, and scanning electron microscopy with energy-dispersive X-ray spectroscopy (SEM–EDS). Following the electron microscopy characterization, samples were imaged by immunofluorescence for the localization of pTau and determination of how this is related to brain cells and the calcification.

## 2. Materials and methods

### 2.1. Samples

Brain tissue from 22 Alzheimer’s disease patients, 22 elderly donors and 15 young (< 40 years old) donors were obtained from the Queen Square Brain Bank and the Brain Tissue Bank of the University of Edinburgh (Supplementary Table S1 has further information about the donors). Tissue samples from the basal ganglia at the level of the anterior commissure, temporal lobe (Brodmann areas 21 and 22), and hippocampal region were selected. Brain tissue samples were formalin fixed, paraffin embedded, cut into 4 µm sections and mounted onto glass histology slides.

### 2.2. Scanning Electron Microscopy

For scanning electron microscopy (SEM) analysis, paraffin wax was removed using pure xylene for two 10-minute intervals. The slides were then mounted on sample holders using double sided carbon adhesive tape, painted with silver conductive paint, and coated with a 5nm carbon layer. A Hitachi S-3499N and a Carl Zeiss LEO 1530 were used at accelerating voltages of 5 kV and 10 kV for SEM imaging, which included secondary electron (SE) and backscattering electron (BSE) modes. Energy dispersive X-ray spectroscopy (EDS) analysis was carried out using Oxford Instruments EDX detectors integrated into both microscopes.

### 2.3. Von Kossa staining

Deparaffinisation and rehydration of samples were done with a sequence of xylene and ethanol washes. Incubation of the samples in a 5% silver nitrate solution (Abcam® ab150687) took place for 45 minutes with 100 Watt incandescent light exposure. The samples were then washed and incubated for 3 minutes in a sodium thiosulphate solution (Abcam® ab150687). Samples then were rinsed with water and incubated in nuclear fast red solution (Sigma-Aldrich N8002) for 5 minutes. Samples were rinsed again with water and dehydrated using absolute ethanol before washing in xylene. A coverslip was placed above the tissue using DPX mounting media (Sigma-Aldrich 06522) and left to dry overnight.

### 2.4. Immunohistochemistry

Deparaffinisation and rehydration of the samples took place through a sequence of xylene and ethanol washes. The samples were then blocked using 1:20 goat serum diluted in a tris buffered saline solution with added 0.1% triton X-100 and 0.5% bovine serum albumin (TBT) for 1 hour, followed by two TBT washes and incubation using a pTau AT8 mouse anti-human primary antibody (Thermofisher MN1020) at a dilution of 1:100. The samples were then washed for 5 minutes three times with TBT and incubated with the secondary antibody (Thermofisher A-21121 and Abcam® ab97239) for an hour at a dilution of 1:200. The samples were then washed and incubated with DAPI (Abcam® ab228549) at a dilution of 1:1000 (diluted in phosphate buffered saline (PBS)) for 15 minutes. The sections were also stained with OsteoSense 680EX (PerkinElmer NEV10020EX) at a dilution of 1:10 (diluted in PBS) for 20 minutes. For mounting, Fluoroshield mounting medium (Abcam® ab104135) was used.

### 2.5. Confocal microscopy

Fluorescence labelling was imaged either using an Olympus FV1000, a Leica SP8 and a Zeiss LSM 980 Airy scan confocal microscopes.

### 2.6. Imaging and colocalization statistical analysis

Image J was used for all image analyses. For colocalization analyses, 10 images per sample were taken using a 63x objective. In total, 10 samples were used: 5 from the elderly donors and 5 from the AD group. For co-occurrence analysis, the Coloc 2 function of Image J was used. All values are reported as median, Interquartile range (IQR). All statistical analyses were done using Origin Lab 2019 and GraphPad Prism 8.3.1 software. A Mann-Witney U test (two-tailed) was used (p<0.05).

## 3. Results and discussion

Von Kossa staining and SEM images clearly show vascular calcification in the brain tissue, in line with findings previously and widely reported in the literature (Supplementary Figure S1). To our surprise, in addition to vascular calcification, both the Von Kossa staining and SEM images of the outer layer (Fig. 1a) of the tissue (the caudate nucleus, the parahippocampal gyrus, and the cortical area of the temporal lobe), show an extended region presenting microcalcifications (Fig. 1b, c, e and f), both in AD patients and in elderly donors (no form of calcification has been observed in young donors).

**Figure 1:**
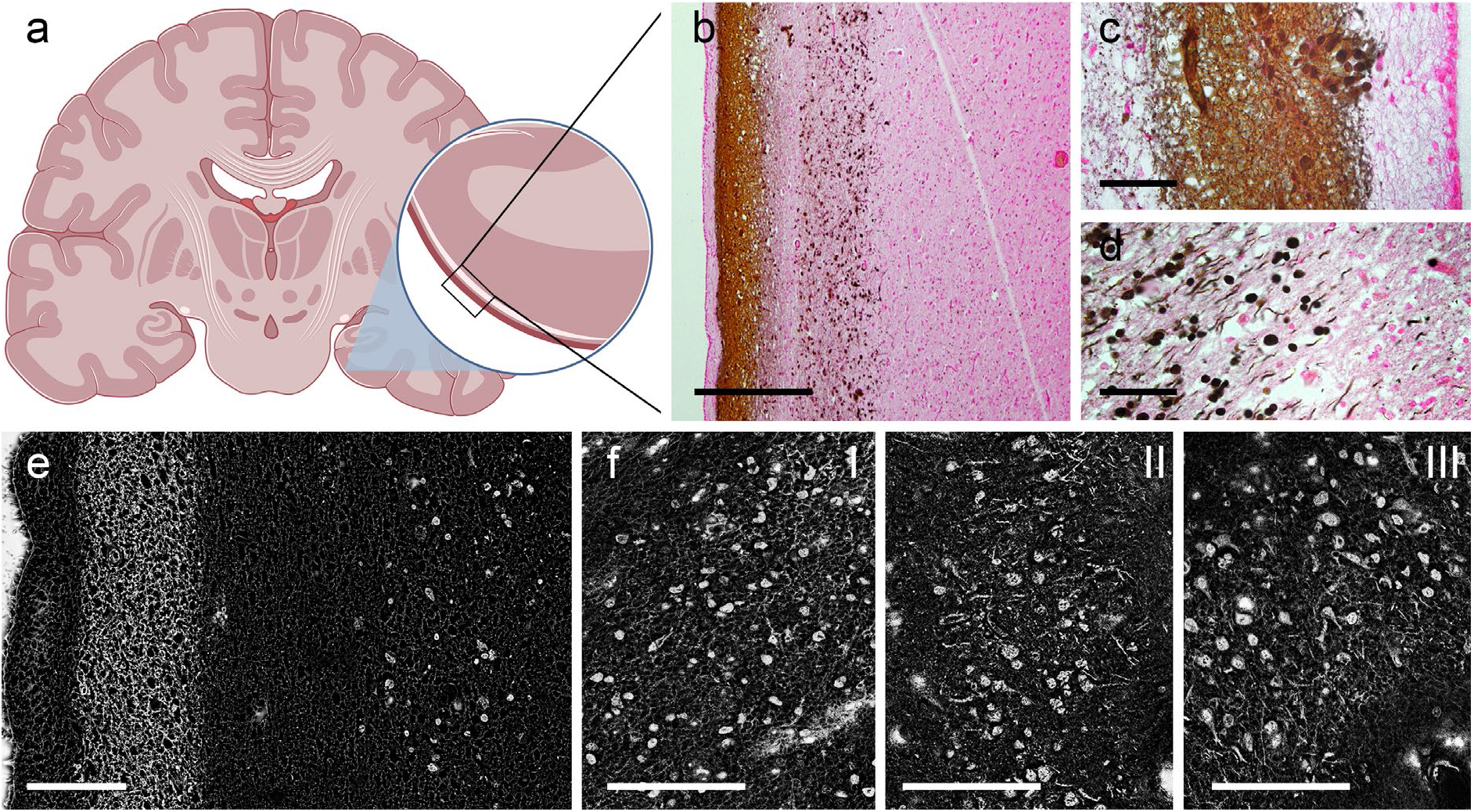
Calcification in histological sections of the human brain. (a) A Human brain diagram indicating one of the outer layers of the tissue where microcalcification was observed. (b) Von Kossa staining of the cortex, revealing microcalcification. Scale bar = 500 µm. (c) Higher magnification image showing von Kossa labelled fibres. (d) Higher magnification image of spherical calcification. Scale bar = 50 µm. (e) Representative backscattered electron micrograph of same region as b. (f) Representative backscattered electron micrograph of basal ganglia I, hippocampus II, and temporal lobe III. Scale bar = 100 µm.

At higher magnification, Density Dependent Colour-SEM (DDC-SEM) micrographs show that the regions with microcalcifications present two main calcified structures: spherical calcification formed from nanoneedles (Fig. 2a and 2c) and calcified fibres (Fig. 2b and 2d), both composed of calcium phosphate, as shown by EDS analysis (Supplementary Fig S2 and S3).

**Figure 2:**
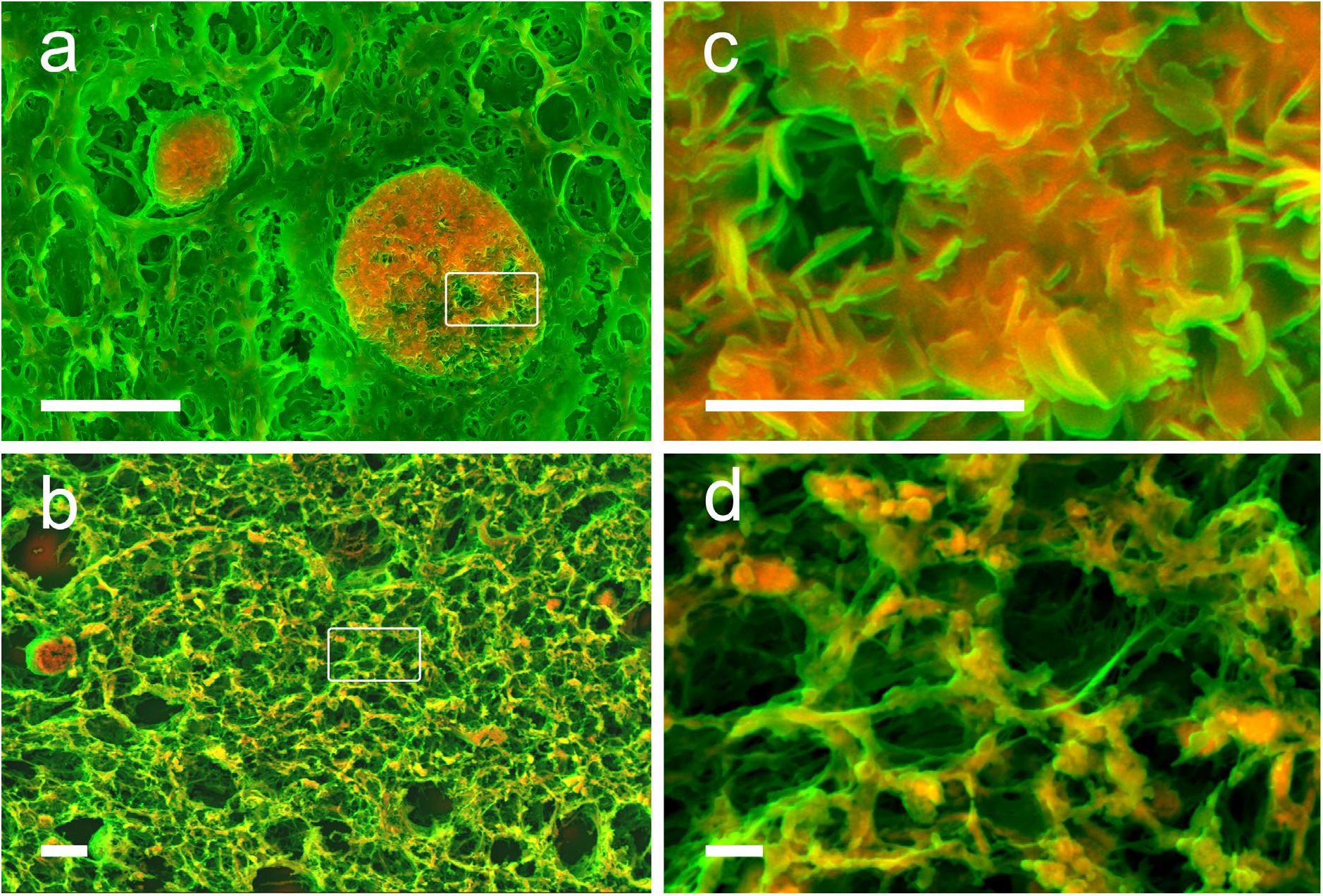
Density Dependent Colour – Scanning Electron Microscopy (DDC-SEM) of calcification in histological sections of the human brain. (a) DDC-SEM of calcified spheres and (b) Calcified fibres. Scale bar = 10 µm. (c) Higher magnification of calcified sphere region highlighted in a, and (d) higher magnification of calcified fibres region highlighted in b. Scale bar = 2 µm.

When comparing the amount of calcification between AD patients and elderly donors, it is clear that AD brains contain a considerably higher amount of calcification (38%, IQR:35.33) than those of elderly donors (4%, IQR:9.515) (Fig. 3a, b and c). Moreover, AD patient samples present more frequently with spherical structures than elderly organ donor samples (Fig. 3d). Interestingly, the size and morphology of the spherical calcification was similar to the cellular nucleus, and some of the calcified fibres are similar to cellular projections, but no calcification observed presented size or morphology similar to the amyloid plaques reported in the literature[34].

**Figure 3:**
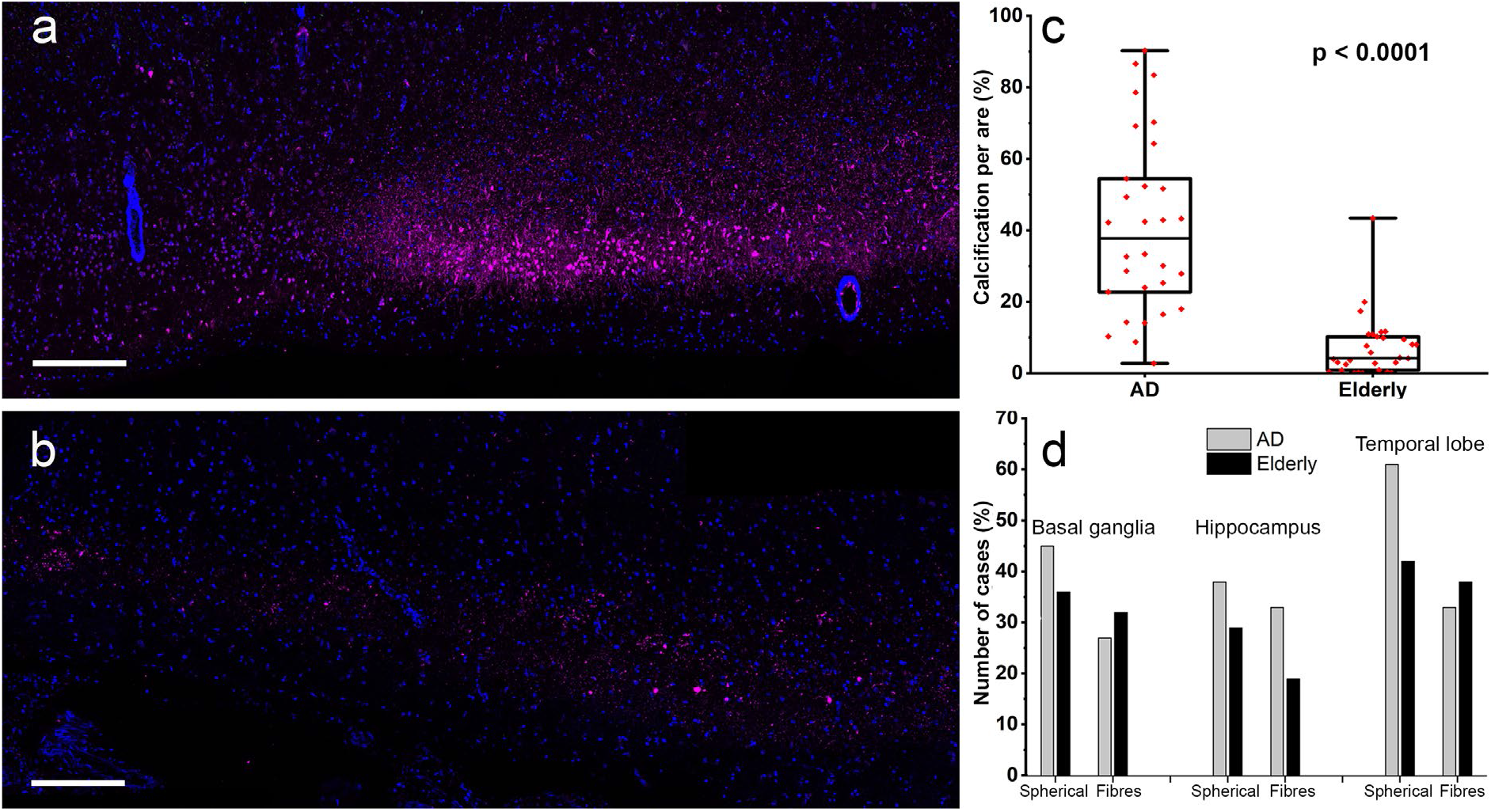
Fluorescence labelling of brain tissue using OsteoSense 680 (magenta) and DAPI (blue). (a) Representative calcification pattern observed in AD brain cases. (b) Representative calcification pattern observed in age matches elderly section. Scale bars = 300 μm. (c) Calcification per area in AD cases and elderly donors (n = 30). Statistical analysis indicated a p-value smaller than 0.0001. Error bars indicate the standard deviation and box lines the upper quartile, median, and lower quartile. (d) Number of patients and organ donors presenting spherical calcification or fibre calcification in different brain regions.

Therefore, our next aim was to investigate whether spherical calcification might be formed in the nucleus of brains cells and whether this calcification might be associated to pTau, since it is well known that this protein accumulates in the nuclei of brain cells[35, 36]. We started off by fluorescently labelling tissue sections from AD patients and elderly donors for DNA, pTau, and calcification.

At lower magnification, fluorescent micrographs show that the staining for calcification closely matches the staining previously observed on SEM micrographs (Fig. 4) of AD patients (Fig.4a and b) and elderly donors (Fig. 4c and d). At higher magnification, fluorescent micrographs show that calcification is indeed present in the nuclei of brain cells (Fig. 5a and b) and other cellular structures, and that this nuclear calcification is significantly higher in AD patients than in elderly organ donors (Fig. 5c). Statistical analysis shows that colocalization of DNA and calcification is 0.78 (IQR: 0.29) in AD patient samples and only 0.06 (IQR: 0.50) in elderly donor samples. The strong colocalization between calcification and DNA in AD patient samples prompts us to suggest that nuclear calcification is strongly associated with AD cases.

**Figure 4:**
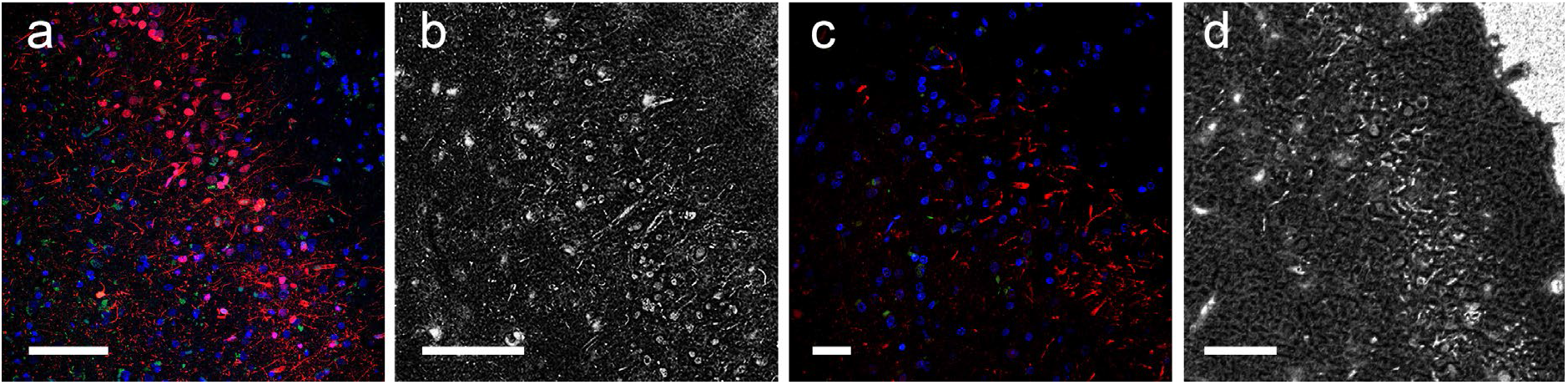
Fluorescence micrographs of hippocampi of AD patients and elderly donors labelled for DNA (blue), pTau (green), and calcification (red). (a) Fluorescence micrograph of the hippocampus interface region from AD brain sample with DNA, pTau, and calcification stains. Scale bar = 77 µm. (b) Scanning electron micrograph of the same region shown in a. Scale bar = 100 µm. (c) Fluorescence micrograph of hippocampus interface region brain sample from elderly organ donor with DNA, pTau and calcification stains. Scale bar = 32 µm. (d) Scanning electron micrograph of the same region shown in c. Scale bar = 50 µm.

**Figure 5:**
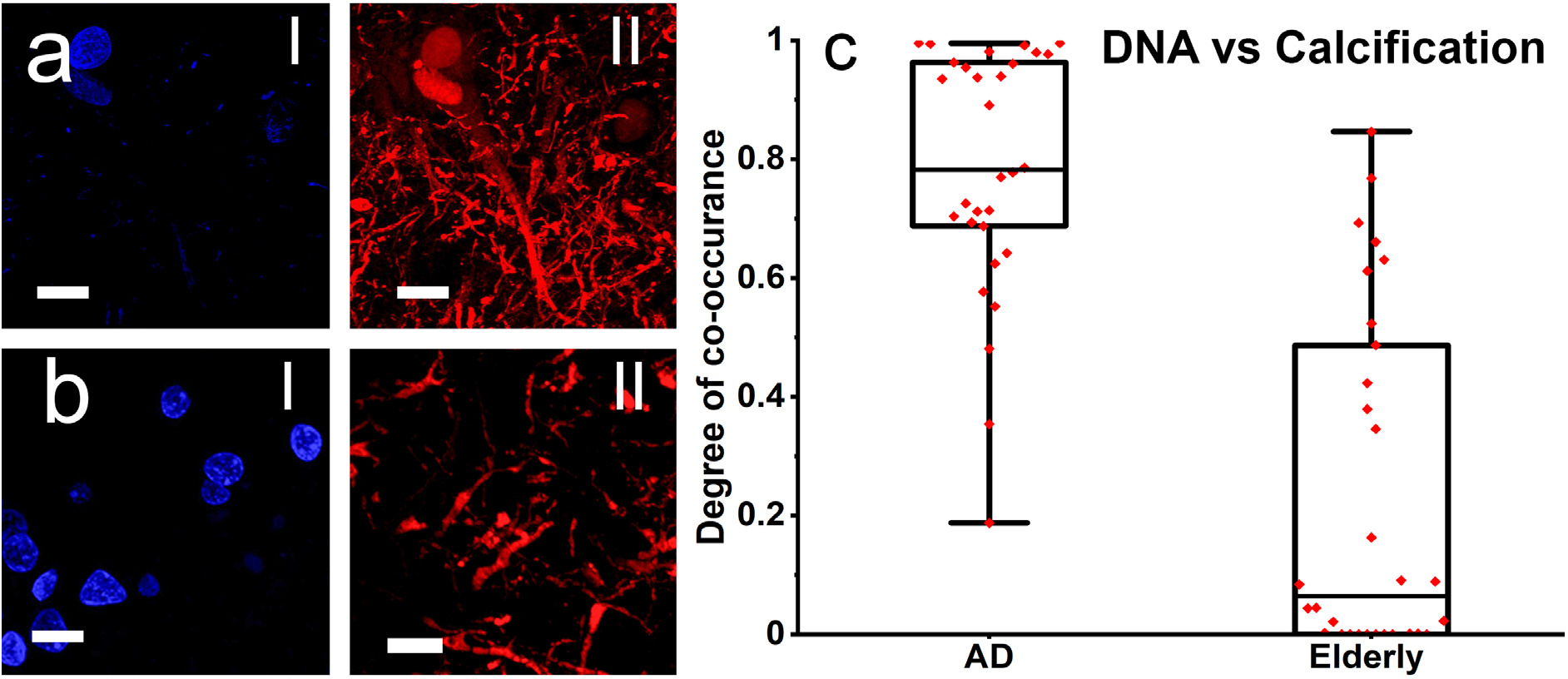
High magnification fluorescence micrographs of AD patients and elderly donors labelled for DNA (blue) and calcification (red). (a) Example of fluorescence micrograph of DNA (I) and calcification (II) from AD patient hippocampus sample. Scale bar = 14 µm. (b) Example of fluorescence micrograph of DNA (I) and calcification (II) from elderly organ donor hippocampus sample. Scale bar = 11 µm. (c) Co-occurrence of DNA and calcification in AD and elderly donors. Wiskers indicate minimum and maximum box lines the upper quartile, median, and lower quartile. p < 0.0001.

In line with previous reports in the literature[35, 36], our fluorescent micrographs also show a correlation between pTau and DNA in AD patient samples, with 0.53 (IQR:0.70) of pTau labelling co-localised with DNA in AD patient samples (Fig. 6a and b) and only 0.12 (IQR: 0.20) in elderly donor samples (Fig. 6c). Finally, fluorescence micrographs (Fig. 7a and b) clearly show that the correlation between pTau and calcification (Fig. 7c) is significantly higher for AD patients samples (0.83, IQR: 0.40) than for elderly organ donors samples (0.19IQR: 0.66).

**Figure 6:**
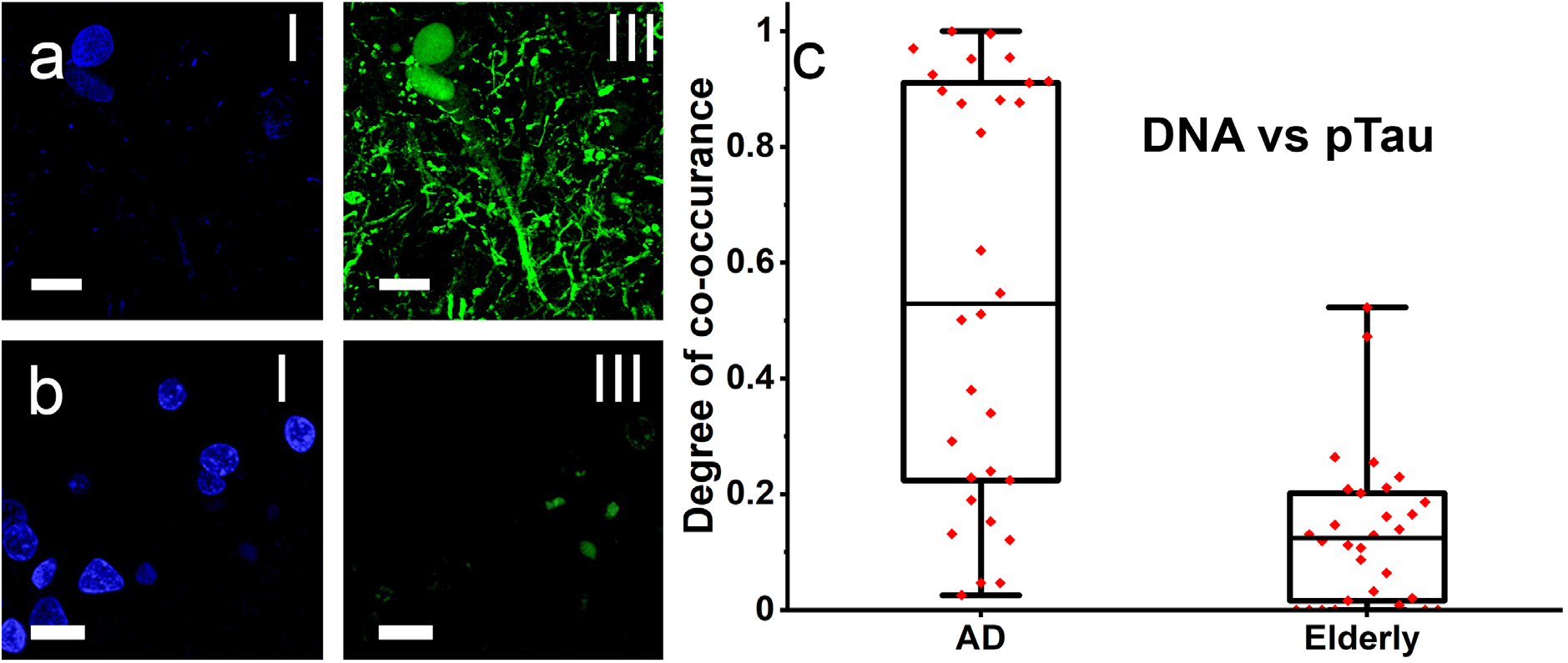
High magnification fluorescence micrographs of AD patients and elderly donors labelled for DNA (blue) and pTau (green). (a) Example of fluorescence micrograph of AD patient hippocampus labelled for DNA(I) and pTau (II). Scale bar = 14 µm. (b) Example of fluorescence micrograph of elderly organ hippocampus labelled for DNA (I) and pTau (II). Scale bar = 11 µm. (c) Co-occurrence of DNA and pTau in AD and elderly donors. Wiskers indicate minimum and maximum box lines the upper quartile, median, and lower quartile. p < 0.0001.

**Figure 7:**
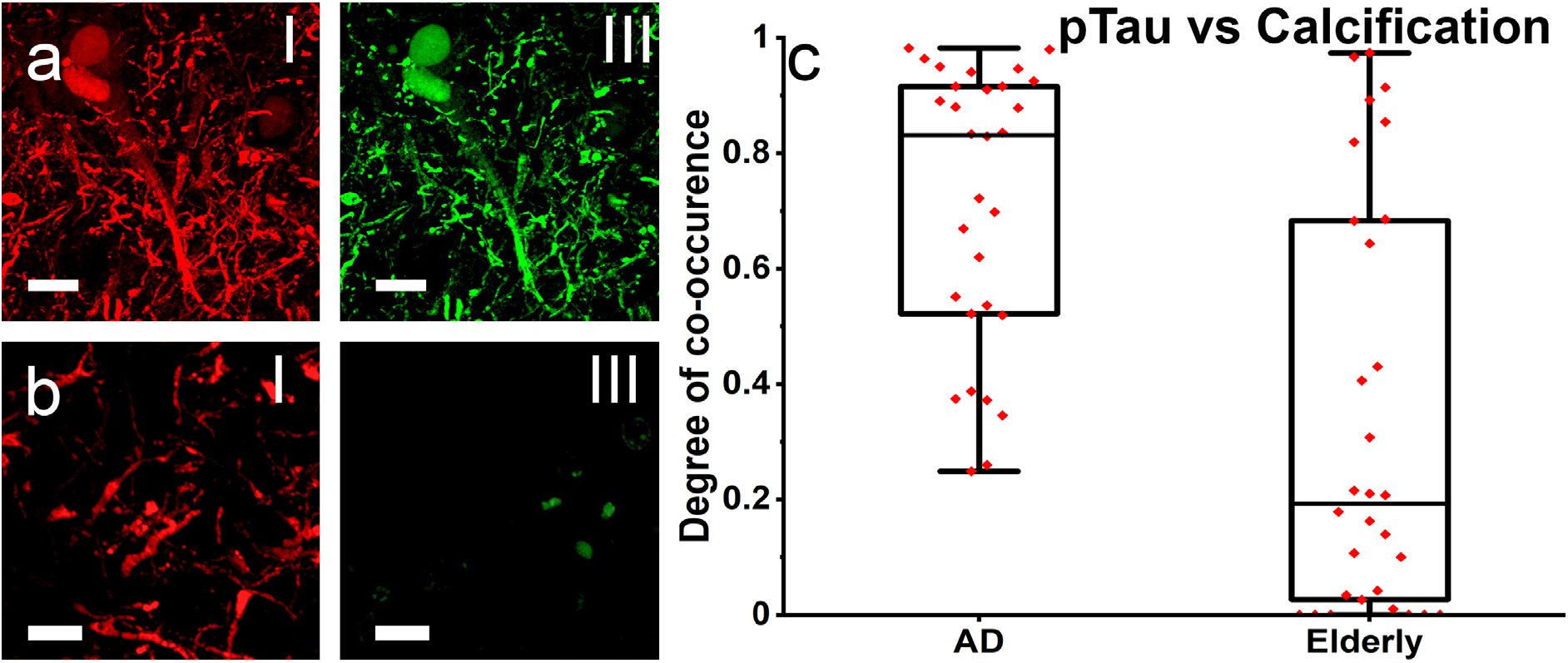
High magnification fluorescence micrographs of AD patients and elderly donors labelled for calcification (red) and pTau (green). (a) Example of fluorescence micrograph of AD patient hippocampus labelled for calcification (I) and pTau (II). Scale bar = 14 µm. (b) Example of fluorescence micrograph of elderly organ donor hippocampus labelled for calcification (I) and pTau (II). Scale bar = 11 µm. (c) Co-occurrence of calcification and pTau in AD and elderly donors. Wiskers indicate minimum and maximum box lines the upper quartile, median, and lower quartile. p < 0.0001.

Taken together, the results presented here show that the calcification of brain cells is strongly associated with pTau presence, particularly with regards to the nuclei of cells in the brains of patients with AD. Nevertheless, it is not yet clear what role calcification plays in the disease. However, a more relevant question should be: which is first to appear, pTau or calcification? Calcification could potentially disturb the metabolism of cells, leading to the hyperphosphorylation of Tau. On the other hand, the process of phosphorylation of Tau (which increases its solubility[37, 38]) could act as a nucleation point for the precipitation of calcium phosphate (by increasing the number of phosphate groups in the protein). Moreover, it is possible that the precipitation of calcium phosphate and Tau phosphorylation could be a synergistic process, which would potentiate both phenomena simultaneously.

The calcification of brain cells might also contribute to explain some intriguing observations from the literature. For instance, take the phenomenon of senescent cells in the brains of AD patients[39], which are no longer functional but are not naturally eliminated from the tissue[40, 41]. Our results suggest that partially or fully calcified cells can still be identified using fluorescent staining and observed by standard microscopy methods. Therefore, it is possible that the odd behaving cells reported in the literature could simply be the calcified cells that may still be stained but are no longer functional.

## 4. Conclusions

The characterization of the calcification in brain cells can contribute to a better understanding of the biochemical mechanisms associated with AD and might also reveal that calcification is part of the full disease mechanism. Calcification could contribute to parallel processes, previously reported in the literature, such as the formation of amyloid plaques and NFT, which would further contribute towards the development and/or progression of AD. Moreover, the association of microcalcification with AD patient brains opens the possibility for calcification to be used as a new biomarker. The calcification can be identified by independent methods, distinct from those currently used for AD diagnosis, assisting in the identification and confirmation of both Tau pathology and AD.

## Acknowledgements

We gratefully acknowledge the roles of the Queen Square Brain Bank and the Brain Tissue Bank of the University of Edinburgh in collecting and making samples and/or data available, as well as of the patients and their next of kin who generously donated tissues and shared data to be used in the research leading to this publication.

## References

[1] R. Brookmeyer, E. Johnson, K. Ziegler-Graham, H.M. Arrighi, Forecasting the global burden of Alzheimer’s disease, Alzheimers Dement. 3(3) (2007) 186–191.

[2] K. Maurer, P. Riederer, H. Beckmann, Alzheimer’s Disease: Epidemiology, Neuropathology, Neurochemistry, and Clinics, Springer-Verlag 1990.

[3] C.A. Lane, J. Hardy, J.M. Schott, Alzheimer’s disease, Eur. J. Neurol. 25(1) (2018) 59–70.

[4] M. Morris, S. Maeda, K. Vossel, L. Mucke, The Many Faces of Tau, Neuron 70(3) (2011) 410–426.

[5] Y. Duan, S. Dong, F. Gu, Y. Hu, Z. Zhao, Advances in the pathogenesis of Alzheimer’s disease: focusing on tau-mediated neurodegeneration, Transl. Neurodegener. 1(1) (2012) 24.

[6] W.H. Stoothoff, G.V. Johnson, Tau phosphorylation: physiological and pathological consequences, Biochim. Biophys. Acta 1739(2-3) (2005) 280–97.

[7] W. Noble, D.P. Hanger, C.C. Miller, S. Lovestone, The importance of tau phosphorylation for neurodegenerative diseases, Front. Neurol. 4 (2013) 83.

[8] R.A. Crowther, Straight and paired helical filaments in Alzheimer disease have a common structural unit, Proc. Natl. Acad. Sci. U S A 88(6) (1991) 2288–92.

[9] L.I. Binder, A.L. Guillozet-Bongaarts, F. Garcia-Sierra, R.W. Berry, Tau, tangles, and Alzheimer’s disease, BBA-Mol. Basis. Dis. 1739(2) (2005) 216–223.

[10] H. Braak, E. Braak, Neuropathological stageing of Alzheimer-related changes, Acta Neuropathol. 82(4) (1991) 239–59.

[11] H. Forstl, A. Burns, N. Cairns, P. Luthert, R. Levy, Basal ganglia mineralization in alzheimers-disease - a comparative-study of clinical, neuroradiological and neuropathological findings, Behav. Neurol. 5(1) (1992) 53–57.

[12] A. Pick, Calcification of the finer cerebral vessels, with remarks upon its clinical significance, Am. J. Psychiat. 61(3) (1905) 417-436-5.

[13] H. Förstl, A. Burns, R. Levy, N. Cairns, Neuropathological Correlates of Psychotic Phenomena in Confirmed Alzheimer’s Disease, Brit. J. Psychiat. 165(1) (1994) 53–59.

[14] A.P.T. Chew, G. Gupta, S. Alatakis, M. Schneider-Kolsky, S.L. Stuckey, Hippocampal Calcification Prevalence at CT: A Retrospective Review, Radiology 265(2) (2012) 504–510.

[15] R. Kockelkoren, J.B. De Vis, W.P.T.M. Mali, J. Hendrikse, P.A. de Jong, A.M. Rozemuller, H.L. Koek, Hippocampal Calcification on Computed Tomography in Relation to Cognitive Decline in Memory Clinic Patients: A Case-Control Study, PLoS One 11(11) (2016) e0167444–e0167444.

[16] R.P. Friedland, J.S. Luxenberg, E. Koss, A Quantitative Study of Intracranial Calcification in Dementia of the Alzheimer Type, Int. Psychogeriatr. 2(1) (1990) 37–43.

[17] D. Bos, M.W. Vernooij, R.F.A.G. de Bruijn, P.J. Koudstaal, A. Hofman, O.H. Franco, A. van der Lugt, M.A. Ikram, Atherosclerotic calcification is related to a higher risk of dementia and cognitive decline, Alzheimers Dement. 11(6) (2015) 639-647.e1.

[18] J. Song, Pineal gland dysfunction in Alzheimer’s disease: relationship with the immune-pineal axis, sleep disturbance, and neurogenesis, Mol. Neurodegener. 14(1) (2019) 28.

[19] J. Wegiel, I. Kuchna, T. Wisniewski, M.J. de Leon, B. Reisberg, T. Pirttila, T. Kivimaki, T. Lehtimaki, Vascular fibrosis and calcification in the hippocampus in aging, Alzheimer disease, and Down syndrome, Acta Neuropathol. 103(4) (2002) 333–43.

[20] S. Bertazzo, E. Gentleman, K.L. Cloyd, A.H. Chester, M.H. Yacoub, M.M. Stevens, Nano-analytical electron microscopy reveals fundamental insights into human cardiovascular tissue calcification, Nat. Mater. 12(6) (2013) 576–583.

[21] S. Bertazzo, E. Gentleman, Aortic valve calcification: a bone of contention, Eur. Heart J. 38(16) (2017) 1189–1193.

[22] E. Tsolaki, W. Doran, L. Magnani, A. Olivo, I.K. Herrmann, S. Bertazzo, Invasive breast tumors are characterized by the presence of crystalline nanoparticles, bioRxiv (2020) 2020.04.29.067660.

[23] A.C. Tan, M.G. Pilgrim, S. Fearn, S. Bertazzo, E. Tsolaki, A.P. Morrell, M. Li, J.D. Messinger, R. Dolz-Marco, J. Lei, Calcified nodules in retinal drusen are associated with disease progression in age-related macular degeneration, Sci. Transl. Med. 10(466) (2018) eaat4544.

[24] D.M.A. Mann, Calcification of the basal ganglia in Down’s syndrome and Alzheimer’s disease, Acta Neuropathol. 76(6) (1988) 595–598.

[25] Y. Iwasaki, M. Ito, K. Mori, A. Deguchi, M. Nagaoka, M. Yoshida, Y. Hashizume, An autopsy case of diffuse neurofibrillary tangles with calcification: early stage pathologic findings, Neuropathology 29(6) (2009) 697–703.

[26] K. Tsuchiya, H. Nakayama, S. Iritani, T. Arai, K. Niizato, C. Haga, M. Matsushita, K. Ikeda, Distribution of basal ganglia lesions in diffuse neurofibrillary tangles with calcification: a clinicopathological study of five autopsy cases, Acta Neuropathol. 103(6) (2002) 555–564.

[27] M.F. Casanova, J.M. Araque, Mineralization of the basal ganglia: implications for neuropsychiatry, pathology and neuroimaging, Psychiat. Res. 121(1) (2003) 59–87.

[28] L. Bäckman, J.L.R. Andersson, L. Nyberg, B. Winblad, A. Nordberg, O. Almkvist, Brain regions associated with episodic retrieval in normal aging and Alzheimer’s disease, Neurology 52(9) (1999) 1861–1861.

[29] G.L. Wenk, Neuropathologic changes in Alzheimer’s disease, J. Clin. Psychiatry. 64 Suppl 9 (2003) 7–10.

[30] R.J. Killiany, M.B. Moss, M.S. Albert, T. Sandor, J. Tieman, F. Jolesz, Temporal Lobe Regions on Magnetic Resonance Imaging Identify Patients With Early Alzheimer’s Disease, Arch. Neurol-Chicago 50(9) (1993) 949–954.

[31] V.L. Villemagne, V. Doré, S.C. Burnham, C.L. Masters, C.C. Rowe, Imaging tau and amyloid-β proteinopathies in Alzheimer disease and other conditions, Nat. Rev. Neurol. 14(4) (2018) 225–236.

[32] R. van der Kant, L.S.B. Goldstein, R. Ossenkoppele, Amyloid-β-independent regulators of tau pathology in Alzheimer disease, Nat. Rev. Neuro. 21(1) (2020) 21–35.

[33] R. Sengoku, Aging and Alzheimer’s disease pathology, Neuropathology 40(1) (2020) 22–29.

[34] M. Querol-Vilaseca, M. Colom-Cadena, J. Pegueroles, R. Nuñez-Llaves, J. Luque-Cabecerans, L. Muñoz-Llahuna, J. Andilla, O. Belbin, T.L. Spires-Jones, E. Gelpi, J. Clarimon, P. Loza-Alvarez, J. Fortea, A. Lleó, Nanoscale structure of amyloid-β plaques in Alzheimer’s disease, Sci. Rep-Uk 9(1) (2019) 5181.

[35] M. Bukar Maina, Y.K. Al-Hilaly, L.C. Serpell, Nuclear Tau and Its Potential Role in Alzheimer’s Disease, Biomolecules 6(1) (2016) 9–9.

[36] J.A. Greenwood, G.V.W. Johnson, Localization and in Situ Phosphorylation State of Nuclear Tau, Exp. Cell Res. 220(2) (1995) 332–337.

[37] N.R. Barthélemy, et al. The Dominantly Inherited Alzheimer, A soluble phosphorylated tau signature links tau, amyloid and the evolution of stages of dominantly inherited Alzheimer’s disease, Nat. Med. 26(3) (2020) 398–407.

[38] N. Sahara, J. Lewis, M. DeTure, E. McGowan, D.W. Dickson, M. Hutton, S.-H. Yen, Assembly of tau in transgenic animals expressing P301L tau: alteration of phosphorylation and solubility, J. Neurochem. 83(6) (2002) 1498–1508.

[39] N. Musi, J.M. Valentine, K.R. Sickora, E. Baeuerle, C.S. Thompson, Q. Shen, M.E. Orr, Tau protein aggregation is associated with cellular senescence in the brain, Aging Cell 17(6) (2018) e12840.

[40] S. Saez-Atienzar, E. Masliah, Cellular senescence and Alzheimer disease: the egg and the chicken scenario, Nat. Rev. Neuro. 21(8) (2020) 433–444.

[41] V. Gorgoulis, P.D. Adams, A. Alimonti, D.C. Bennett, O. Bischof, C. Bishop, J. Campisi, M. Collado, K. Evangelou, G. Ferbeyre, J. Gil, E. Hara, V. Krizhanovsky, D. Jurk, A.B. Maier, M. Narita, L. Niedernhofer, J.F. Passos, P.D. Robbins, C.A. Schmitt, J. Sedivy, K. Vougas, T. von Zglinicki, D. Zhou, M. Serrano, M. Demaria, Cellular Senescence: Defining a Path Forward, Cell 179(4) (2019) 813–827.

