## Supplementary materials for "Nuclear and cellular, micro and nano calcification in Alzheimer’s disease patients and correlation to phosphorylated Tau"

### Supplementary material

**Table S1: Clinical information from donors of samples prepared in the form of histological slides.**

| Case | Sex | Patient Age | Clinical information | Category |
|---|---|---|---|---|
| 1 | M | 64 | Braak V1, CERAD definite, moderate CAA, AD diagnosis | AD |
| 2 | F | 56 | Braak VI,AD diagnosis | AD |
| 3 | F | 62 | Braak VI, severe CAA, limbic TDP-43 pathology, amygdala predominant LB pathology, AD diagnosis | AD |
| 4 | M | 75 | moderate CAA, LB pathology, AD diagnosis | AD |
| 5 | F | 79 | Braak VI, moderate CAA, mild hyaline arteriosclerosis, mild atherosclerosis, AD diagnosis | AD |
| 6 | M | 80 | Braak V, transitional LB pathology, mild CAA, mild small vessel disease, warershed infacrtion in frontal lobe, AD diagnosis | AD |
| 7 | M | 77 | severe CAA, transitional LBD, AD diagnosis | AD |
| 8 | F | 88 | Braak V, severe CAA, CVD of small vessel type, AD diagnosis | AD |
| 9 | M | 57 | AD diagnosis | AD |
| 10 | M | 76 | Braak V, severe CAA, AD diagnosis | AD |
| 11 | F | 67 | Braak VI, severe CAA, AD diagnosis | AD |
| 12 | F | 71 | Moderate CAA, AD diagnosis | AD |
| 13 | F | 79 | Braak VI, moderate small vessel disease, AD diagnosis | AD |
| 14 | M | 69 | severe CAA, diffuse white matter injury associated with severe CAA, AD diagnosis | AD |
| 15 | F | 76 | AD diagnosis | AD |
| 16 | F | 61 | Braak VI, Moderate CAA, lacunar infarct, AD diagnosis | AD |
| 17 | F | 76 | Braak VI, severe CAA, TDP-43 proteinopathy, AD diagnosis | AD |
| 18 | M | 62 | severe CAA, AD diagnosis | AD |
| 19 | F | 66 | Braak VI, hippocampal sclerosis, AD diagnosis | AD |
| 20 | M | 72 | Braak VI, TDP-43 proteinopathy, AD diagnosis | AD |
| 21 | M | 68 | Braak VI, AD diagnosis | AD |

| | | | | |
|---|---|---|---|---|
| 22 | M | 88 | AD diagnosis | AD |
| 23 | F | 82 | Pathological aging Braak II, mild CAA | Elderly |
| 24 | F | 56 | mild small vessel disease | Elderly |
| 25 | F | 87 | mild age related changes (Braak I), mild CVD | Elderly |
| 26 | F | 88 | Normal, multiple cortical infarcts, small vessel disease and pathological ageing | Elderly |
| 27 | F | 78 | | Elderly |
| 28 | M | 95 | lacunar infarct due to small vessel disease, pathological ageing (Braak 1) Argyrophillic grain disease | Elderly |
| 29 | M | 85 | | Elderly |
| 30 | F | 80 | Argyrophilic brain disease, pathological ageing Braak II, small vessel disease | Elderly |
| 31 | M | 83 | Pathological ageing (Braak III) | Elderly |
| 32 | F | 79 | Pathological ageing Braak 1, mild hyaline arteriosclerosis | Elderly |
| 33 | F | 86 | Small vessel disease | Elderly |
| 34 | M | 87 | moderate small vessel disease, tau pathology (Braak I) | Elderly |
| 35 | F | 82 | Pathological ageing (Braak II), mild small vessel disease, moderate CAA | Elderly |
| 36 | M | 87 | Pathological ageing (Braak II) | Elderly |
| 37 | F | 89 | Vascular dementia, old cerebral infarct, AD path low | Elderly |
| 38 | F | 78 | mild small vessel disease, mild age-related changes | Elderly |
| 39 | F | 68 | metastasis | Elderly |
| 40 | F | 87 | path aging Braak II, mild CVD | Elderly |
| 41 | F | 80 | argyrophilic brain disease, pathological ageing Braak II, small vessel disease | Elderly |
| 42 | M | 88 | | Elderly |
| 43 | F | 93 | pathological ageing (Braak III) | Elderly |
| 44 | F | 92 | cerebral infarct, CVD, pathological ageing (Braak III) | Elderly |
| 45 | M | 17 | Severe head and neck trauma, 1b - Road traffic collision (motorcycle rider), Brain Diffuse vascular trauma | Young |
| 46 | M | 25 | Extensive Internal Haemorrhage,1b - Multiple Injuries,1c - Road traffic collision (driver) | Young |
| 47 | F | 20 | Suspension by a ligature | Young |
| 48 | F | 40 | | Young |
| 49 | M | 40 | | Young |
| 50 | M | 39 | Suspension by a ligature | Young |
| 51 | M | 34 | | Young |
| 52 | M | 26 | Multiple injuries owing to a road traffic accident | Young |
| 53 | M | 19 | | Young |
| 54 | M | 25 | Multiple Injuries,1b - Road traffic collision (passenger) | Young |
| 55 | M | 25 | Internal bleeding,b - Rupture of the thoracic aorta, | Young |

| | | | | |
|---|---|---|---|---|
| | | | 1c - Road traffic collision (driver), Possible grade 3 diffuse axonal injury (DAI). | |
| 56 | M | 21 | Stab wound to chest | Young |
| 57 | M | 22 | PM toxicology - hypoglycaemia. | Young |
| 58 | M | 16 | Suspension by a ligature | Young |
| 59 | M | 28 | Suspension by a ligature | Young |

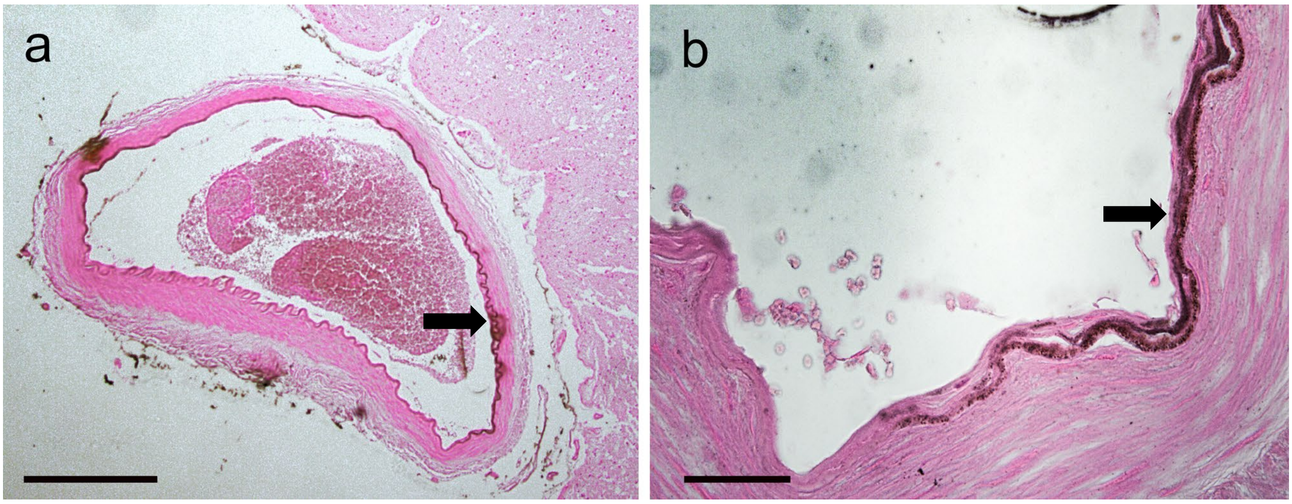


**Figure S1:** Tissue calcification as observed using von Kossa staining in the basal ganglia of elderly donors and AD patients. a Low magnification optical micrograph of vascular calcification in the basal ganglia of an AD patient. Scale bar = 500 μm. **b** High magnification optical micrograph of vascular calcification in the basal ganglia of an AD patient. Scale bar = 50 μm.

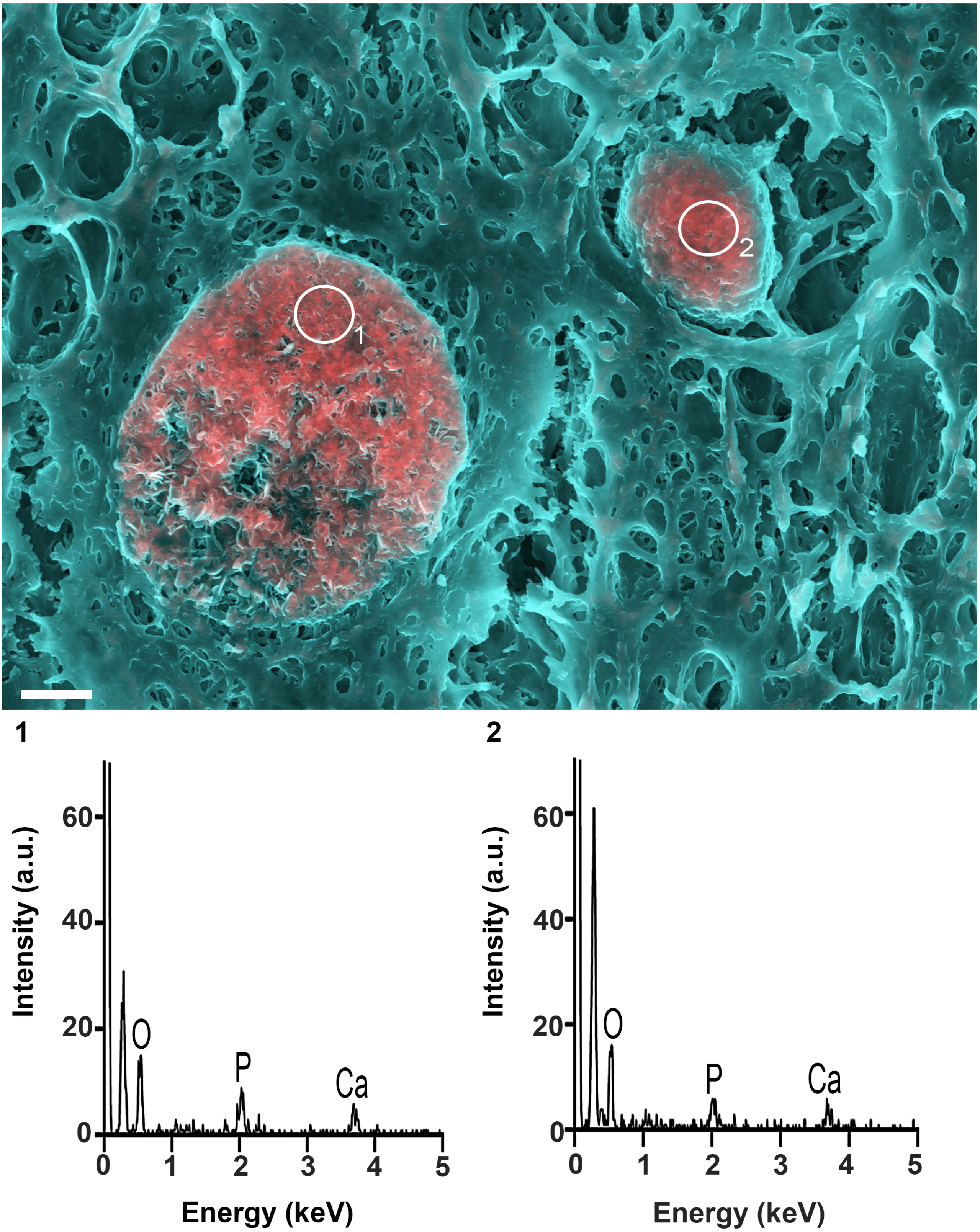


**Figure S2:** DDC-SEM image of calcified spheres observed in the basal ganglia, where pink/red indicate calcification and turquoise indicates tissue. EDS analysis indicated that all calcified spheres, regardless of size, were composed of calcium and phosphorus. Scale bar = 2 µm.

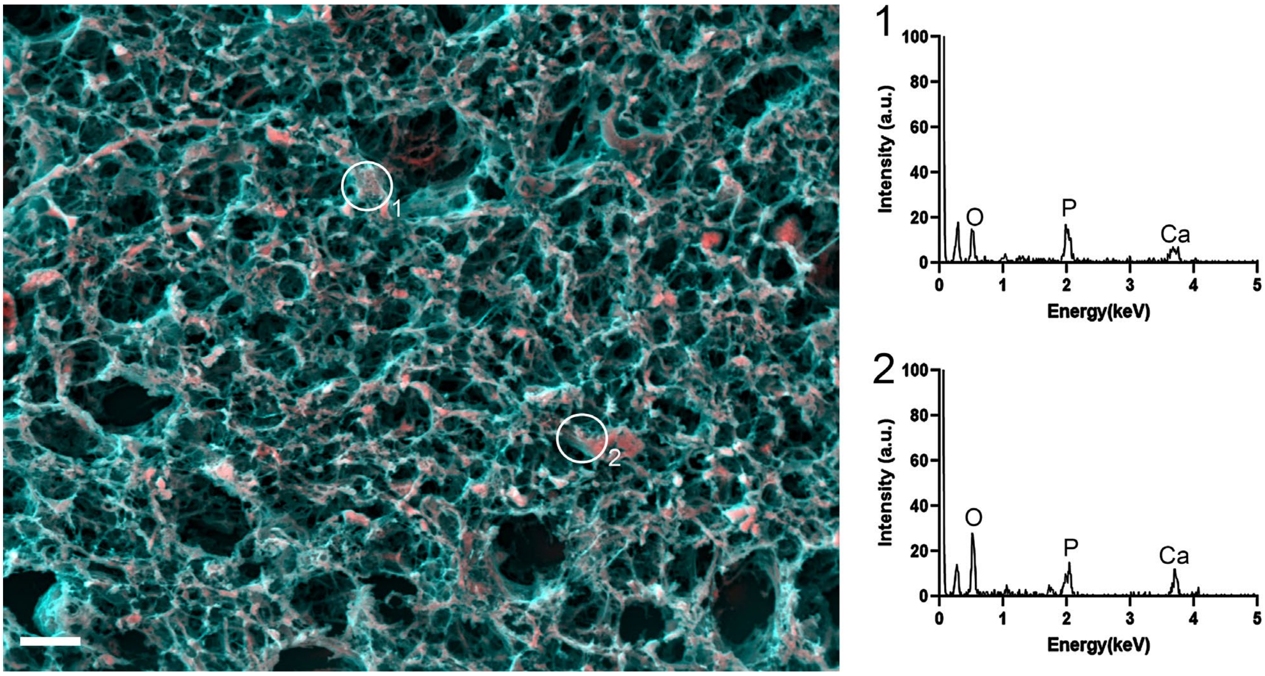


**Figure S3:** DDC-SEM image of calcified fibres observed in the basal ganglia where pink/red indicates calcification and turquoise indicates tissue. EDS analysis indicated that all calcified fibres were composed of calcium and phosphorus. Scale bar = 10 µm.
